# Bilateral field advantage of spatial attention in macaque lateral prefrontal cortex

**DOI:** 10.1101/2024.04.29.591631

**Authors:** Maryam Nouri Kadijani, Theda Backen, Kaustubh Machanda, Sandeep K Mody, Stefan Treue, Julio C. Martinez-Trujillo

## Abstract

Allocating visual attention to behaviorally relevant stimuli is easier when distractors are located in the opposite visual hemifield relative to when they are in the same hemifield. The neural mechanisms underlying this bilateral field advantage remains unclear. We documented this effect in two macaques performing a covert spatial attention task in two different conditions: when the target and distracter were positioned in different hemifields (across condition), and when they were positioned on the top and bottom quadrants within the same visual hemifield (within condition). The animals’ behavioral performance at detecting a change in the attended stimulus was higher in the across relative to the within condition. We recorded the responses of lateral prefrontal cortex (LPFC, area 8A) neurons in one animal. The proportion of LPFC neurons encoding the allocation of attention was larger in the across relative to the within condition. The latter was accompanied by an increase in the ability of single neurons to discriminate the allocation of attention in the across relative to the within condition. Finally, we used linear classifiers to decode the allocation of attention from the activity of neuronal ensembles and found a similar bilateral field advantage in decoding performance in the across relative to the within condition. Our finding provides a neural correlate of the bilateral field advantage reported in behavioral studies of attention and suggest that the effect may originate within the LPFC circuitry.

## INTRODUCTION

Visual attention prioritizes the processing of behaviorally relevant visual stimuli while filtering out distracters (Desimone & Duncan, 1995). Studies in primates have shown that attending to a stimulus at one location selectively enhances the responses of neurons with receptive fields (RFs) at that location while relatively suppresses responses of neurons with RF at other locations (Desimone & Duncan, 1995; Maunsell, 2015; Moran & Desimone, 1985; Reynolds et al., 1999; Treue & Martinez-Trujillo, 1999; Treue & Maunsell, 1996). Remarkably previous studies have shown that the behavioral benefits of spatial attention are more pronounced when targets and distractors are presented in different visual hemifields compared to when presented within the same hemifield (Alvarez & Cavanagh, 2005; Cavanagh & Alvarez, 2005; Chakravarthi & Cavanagh, 2009; Delvenne, 2005). The neurophysiological bases of this bilateral field advantage of attention remain poorly investigated at the level of single neurons and cortical microcircuits.

Previous studies in human subjects (Störmer et al., 2014; Vandenberghe et al., 2000) have suggested that there are independent processing resources for contralateral and ipsilateral visual hemifields. Using Event related potentials (ERPs) recordings the latter study found that early visual cortical areas show stronger correlates of the bilateral field advantage effect compared to association areas. Matsushima and Tanaka (2014) performed single-unit recordings in the lateral prefrontal cortex (LPFC) of macaques while animals held in working memory the spatial location of one or two stimuli (Matsushima & Tanaka, 2014). They found reduced neuronal activity when animals remembered the locations of two stimuli within the same hemifield as compared to when they remembered stimuli distributed across the visual hemifields. Buschman and colleagues (2011) recorded single units from macaque LPFC also during a working memory task and reported decreased behavioral performance as well as impaired information coding when distracters were in the same hemifield as the target relative to when they were in opposite hemifields (Buschman et al., 2011). Thus, at the resolution of single neurons areas downstream from early visual areas show correlates of the bilateral field advantage.

One possibility is that the computations underlying attentional filtering of stimuli distributed across the visual field occur in areas downstream from early visual cortex where neurons have a bilateral representation of the visual field (Bullock et al., 2017). This may facilitate microcircuit computations between subpopulations of neurons within a local network. One of such areas is the lateral prefrontal cortex (LPFC). Some studies in macaque LPFC, specifically areas 8/9/46, have previously demonstrated that neurons show strong response modulation with attention (Bichot et al., 2015; Duong et al., 2019; Everling et al., 2002), that such a modulation is correlated with the behavioral performance of the animals (Lennert & Martinez-Trujillo, 2011), and occurs earlier in LPFC than in upstream visual areas (Lennert & Martinez-Trujillo, 2013). This suggests that neural activity in the LPFC is causally related to the effects attention on information processing in the brain. Moreover, different from early visual areas, each LPFC has a bilateral representation of the visual field (Bullock et al., 2017; Duong et al., 2019; Tremblay et al., 2015), which can facilitate dynamic competitive interactions between neurons representing stimuli within the same as well as across different hemifields (Desimone & Duncan, 1995; Duong et al., 2019). One possible scenario is that competitive interactions during the allocation of attention are less intense when distracters are in the opposite hemifield as the target compared to distracters within the same hemifield. The latter would allow better representation of target features when distracters fall in the opposite hemifield.

To test this hypothesis, we simultaneously recorded the activity of neurons in area 8A, which is a cytoarchitectonically defined granular region of the LPFC, of two macaque monkeys by using a 10×10 multielectrode array (Utah array, (Normann & Fernandez, 2016)). We recorded activity of single neurons and ensembles while the animals covertly allocated attention to one stimulus (the target) and ignored a second stimulus (the distractor). The stimuli were presented one to the left and the other to the right of a fixation point (across hemifields) or both on the same side relative to the fixation point (within hemifield). Consistent with later behavioral studies, we found a behavioral performance was higher in the across than in the within hemifield condition. The proportion of single units showing selectivity for attended/unattended stimuli was higher in the across than in the within condition. From the neural ensemble activity, we decoded the focus of attention in both tasks with performances significantly higher than predicted by chance. Decoding performance was higher in the across relative to the within condition.

## MATERIALS & METHODS

### Animals

Two adult male rhesus monkeys (*Macaca mulatta,* R: 9.7 kg; S: 10.2 kg) participated in the experiments. All animal procedures complied accordance with McGill University’s animal care committee’s regulations. During the training and testing periods, the animals received their minimum daily amounts of fluids so that they were motivated to perform the tasks. Animal body weight was monitored during the duration of the experiments to make sure non weight loss or deterioration in the animals’s health happened.

### Visual Stimuli

The stimuli were back projected on a screen using a video projector (NEC WT610, 1024 x 768 pixel resolution, 75 Hz) and custom-made software running on an Apple G4 Power PC. The animals viewed the screen at a distance of 57 cm (1 cm = 1° of visual angle). The stimuli were generated by plotting colored dots (white, 76.39 cd/m2; grey, 10.83 cd/m2; pink, 22.68 cd/m2; green, 11.26 cd/m2; blue, 10.96 cd/m2; red, 8.92 cd/m2; turquoise, 44.14 cd/m2) on a dark background (black-gray, 0.74 cd/m2) with a density of three dots per degree^2^ within a circular stationary virtual aperture. All dots within one random dot pattern (RDP) moved coherently at a speed of 15°/s and were replotted at the opposite side when they crossed the border of the aperture. The radius of the aperture was 4°. In the across-hemifield condition, the RDPs’ centers were 8° away from the fixation spot and the two stimuli were 16° apart, whereas in the Within-hemifield condition, their centers were also presented at 8° eccentricity but on the diagonal, which made the two stimuli 11.33° apart. We chose the differences in distance between the stimuli to keep eccentricity constant, which influenced the animals’ performance when designing the experiments.

### Task

The animals initiated a trial by keeping gaze within a 2° diameter window (4° in monkey S) centered on a small fixation spot (0.24^(^°^)2^). Gaze position was monitored using an infrared video-based eye tracker (EyeLink 1000, SR Research, Ontario, Canada). After a 353 ms fixation period, two moving RDPs appeared at two different locations. The position of the RDPs depended on the condition, either the left or right of the fixation point (across condition) or both in the same side of the fixation point (within condition). The patterns were composed of white dots on a dark background that moved either up (0°) or down (180°) relative to the vertical. After a variable interval (294, 471, or 647 ms) following the RDPs’ onset, both patterns changed to different colors (e.g., one to green and the other to red). The task for the animals was to identify one of the two RDPs as the target based on its color and covertly attend to it while ignoring the other (the distractor). After 706 ms, the color was removed, and the RDPs returned to white. The animals had to maintain attention on the target and wait between 753 ms to 1600 ms for a brief motion direction change in the target stimulus (118 ms duration, change was 32° clockwise from the current direction) and release the button within 100 ms-650 ms. In 50% of the trials, the distractor changed motion direction before the target. In those trials, the monkey had to keep holding the lever until the target changed. Which of the two colors indicated the target was based on an ordinal color-rank rule the monkey had learned over the training sessions (turquoise > red > blue > green > pink > grey, see also Lennert & Martinez-Trujillo 2011). Each correctly performed trial was rewarded with a drop of juice. A sequence of correct trials yielded a slight increase in reward size. Trials in which the monkey responded to the distractor change (false alarms) or did not respond to the target change within the reaction time window (misses) or broke fixation before the end of a trial (fixation breaks), were terminated without reward. The different trial types were presented in random sequence. Only correctly, performed trials were included in the analysis unless otherwise indicated.

### Surgical Procedures

The surgical operations were carried out under general anesthesia using isoflurane administered through endotracheal intubation. The animals were implanted with titanium head posts used to restrain head motion. We chronically implanted a 10×10 multi-electrode array (96 channels, Blackrock Microsystems LLC, Utah, USA) in each monkey’s left LPFC. The array was positioned on the cortical surface anterior to the knee of the arcuate sulcus and caudal to the posterior end of the principal sulcus, known as area 8a in the macaque monkey (Petrides et al., 2012).

### Electrophysiological Recordings

We recorded from all 96 active channels (out of the 10×10 = 100 channels, there are four reference channels in the MEA) from the left LPFC of both animals. Data were recorded using a Cerebus Neuronal Signal Processor (Blackrock Microsystems LLC, Utah, USA) via a Cereport adapter. After 1x amplification in the head stage (ICS-96), the neuronal signal was band-pass filtered (0.3 Hz-7.5 kHz) and digitized (16) at a sample rate of 30 kHz. For each channel, spike waveforms were detected by manually thresholding (∼4 times the root mean square of the noise amplitude) the digitally high-pass filtered (250 Hz/4 pole) raw voltage trace. The extracted spikes and associated waveforms were sorted offline using both manual and semi-automatic techniques using OfflineSorter (Plexon Inc, TX, USA) and Matlab (MathWorks, Natick, MA, USA). We made no assumptions as to whether the recorded units were same or different from day to day.

### Data Analysis

Analysis of spike data (firing rates) and statistical tests were performed using MATLAB. Our analyses were computed for different epochs. A 350 ms window during fixation period and a 500 ms window during baseline, cue (150 ms to 650 ms after color onset) and post-cue/sustained attention (200 ms to 700 ms after color offset) (see **Fig. 2**).

**Figure 1:**
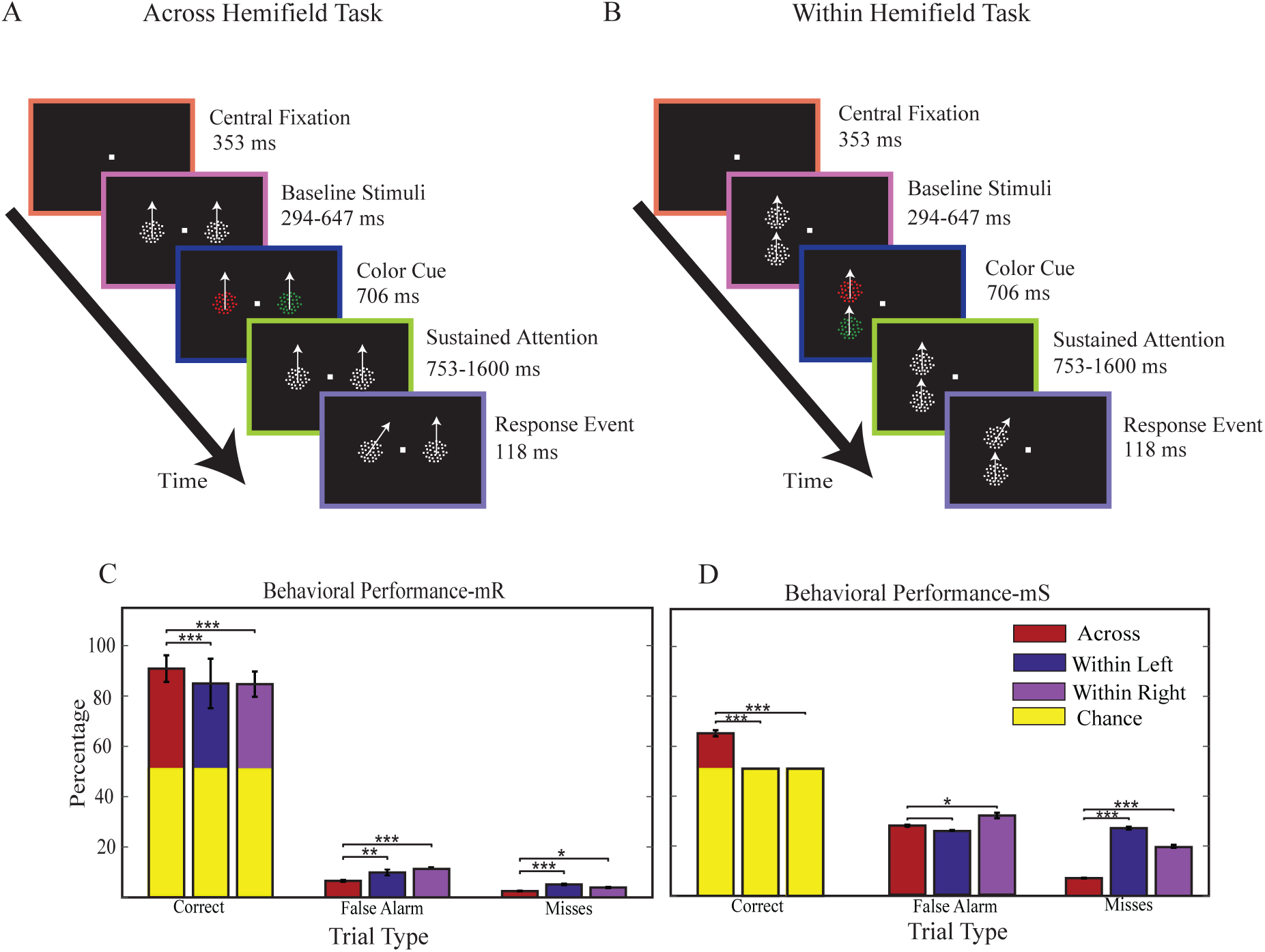
Behavioral Task and Performance. (**A**) Example trial of the across task with stimuli distributed the left and right visual fields. (**B**) Example trial of the within task with both stimuli in the ipsilateral of recording side (left hemifield). (**C-D**) Percentage of correct, false and miss trials for the across task (red bars), the within left task (blue bars) and the within right task (magenta bars) for both monkeys (**C**: monkey R, **D**: monkey S). Error bars represent standard deviation across sessions. Asterisks mark significant differences in mean performance (*χ*^2^ test; * p < 0.05, ** p < 10^-2^ and *** p < 10^-3^).

**Figure 2:**
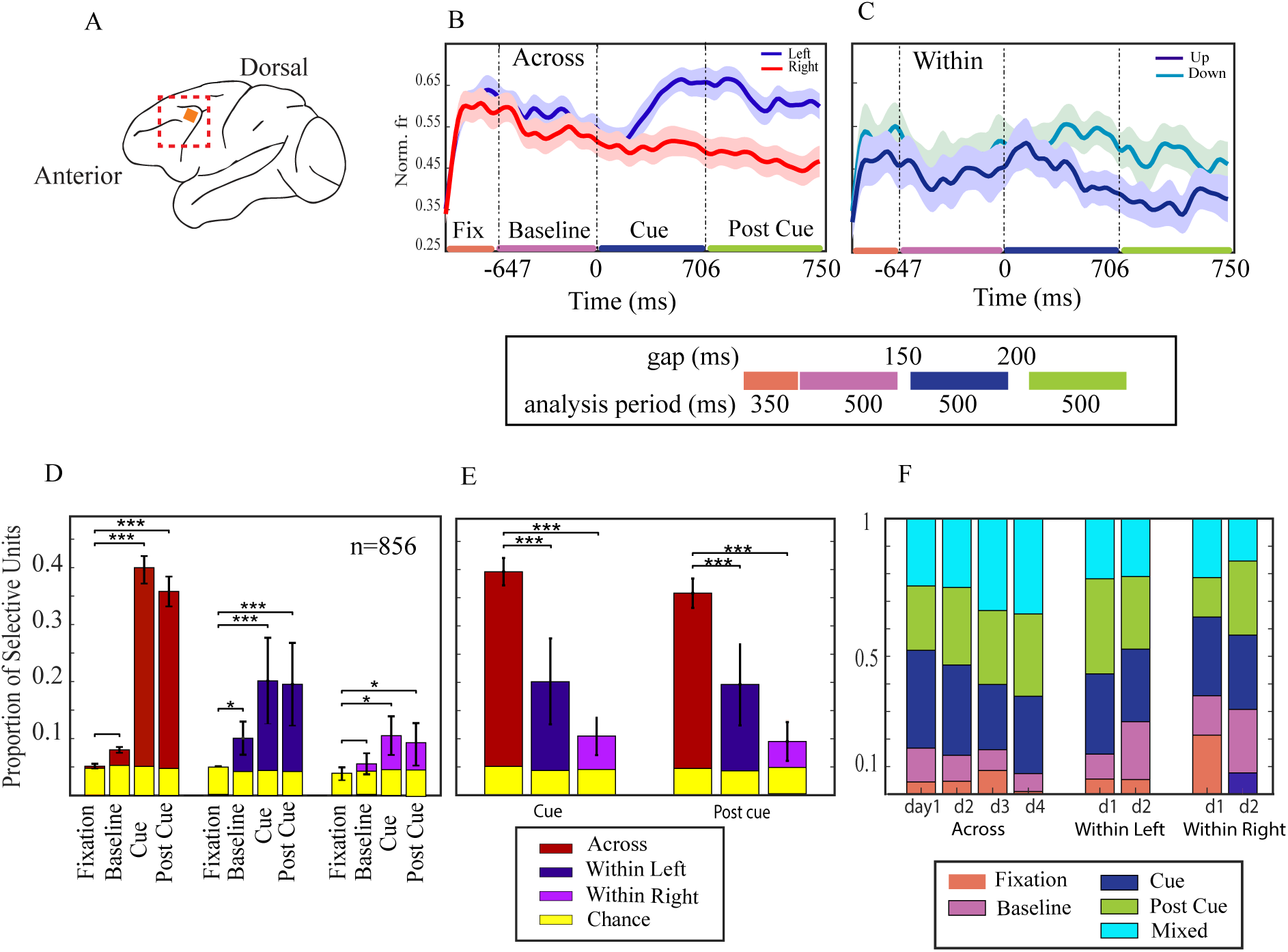
Array Location and Electrophysiological Recording. **A**) Schematic of a macaque brain with area 8a of LPFC highlighted and anatomical location of the multielectrode array implanted in monkey R (top). **B, C**) Mean Normalized responses of tuned units to the different target positions from monkey R recorded during **(B)** the across condition (ipsilateral target: light blue, contralateral target: light red) and **(C)** the within condition (target up: light blue, target bottom: dark blue) as a function of time. Shading represents SEM (±) at each time point. Colored horizontal lines represent different time epochs of experiment: fixation (orange), baseline (magenta), cue (dark blue) and post cue (light green) which are consistent in all figures. (**D**) Proportion of location-selective units (one-way ANOVA with factor target location, p < 0.05) for four different condition periods in the across condition (red bars), the within left (blue bars), the within right (magenta bars) and chance level (yellow bars). Error bars represents SEM (±) over recording sessions. Asterisks mark significant differences in mean proportion (χ^2^-test; * p < 0.05, ** p < 0.01, *** p < 0.001). (**E**) Proportion of tuned units in cue and post cue epochs across three conditions. (**F**) Proportion of tuned units in each recording sessions to see whether the distributions of tuned units were approximately stable over time. The colors indicate trial periods.

**Figure 3:**
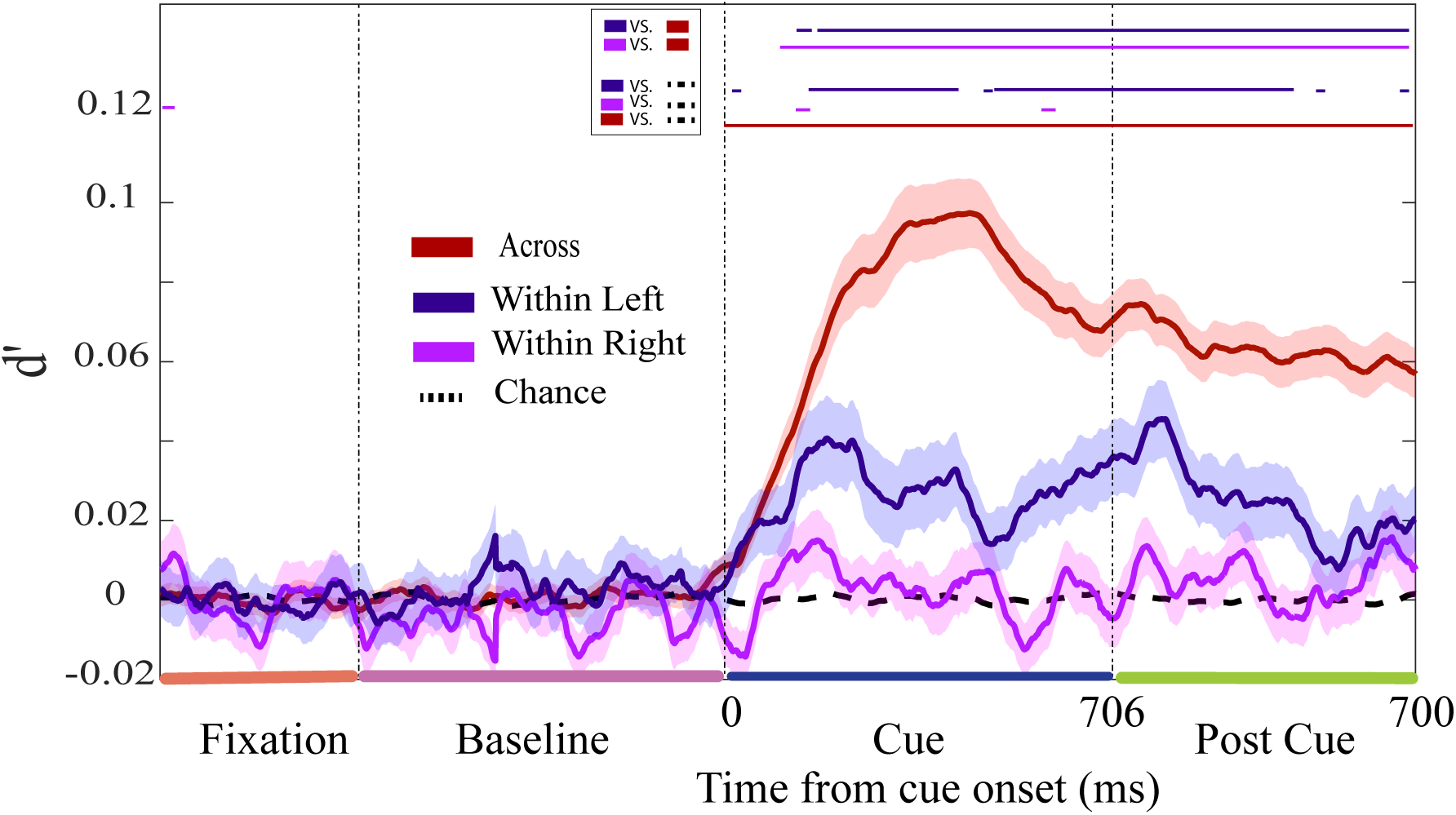
Neuronal discriminability. Average d’ for discriminating the allocation of attention as a function of time for selective units recorded during across (red), within left (dark blue) and within right conditions (magenta). Shading represent SEM (±). Horizontal lines on top indicate at which time bins d’ was significantly different from chance level and across condition (Wilcoxon rank-sum test, p < 0.05).

We recorded the response of 856 neurons in monkey R (525 over 4 sessions of the across condition and 331 over 4 sessions of the within condition). We excluded neurons with a firing rate less than Hz.

### Neuronal Selectivity

We determined neuronal selectivity by performing a one-way analysis of variance (ANOVA) with target location as a factor using mean activity of each neuron within a 100 ms window starting from each epoch onset and slid along the trial in steps of 10 ms independently for four different epochs. If the neuron revealed a significant effect of target position (p< 0.05) in at least five consecutive time bins, it was classified as a target selective unit. The position of the target (left or right in across condition and up or down in within conditions) at which the unit produced stronger response (two sample t-test p<0.05) was considered the neuron’s preferred location (Lennert & Martinez-Trujillo, 2011, 2013).

We calculated the proportions of location-selective cells in the entire population in the different periods for each task. We compared the proportions between periods and conditions using a Chi-Square test (χ^2^, p < 0.05). In order to determine whether these proportions were different from those expected by chance, we compared them to those obtained using a randomization procedure (chance level). For the randomization procedure, we used the same trials and units as in the original data but shuffled the trial orders.

### Spike Density Function

The activity of selective neurons were plotted as mean spike density functions (SDF) over trials, generated by convolving the spike train with a Gamma kernel function (growth parameter 1 ms and decay parameter 20 ms) (Lennert & Martinez-Trujillo, 2013). We normalized SDF for each neuron by the maximum firing rate of the neuron.

### Neuronal Sensitivity

To quantify the ability of single neurons to discriminate between targets and distractors, we used d’, which was calculated as follows:

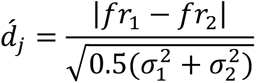

where ‘*d_j_* is sensitivity index for neuron j, *fr_i_* is mean response of neuron j to the two possible target locations i (i=1,2) and *σ_j_* is variance of the responses over target location i. The d’ values were calculated for a 100 ms window slid in steps of 1 ms along the spike train for each selective neuron. We also shuffled the trial order 50 times for each neuron and computed d’ to obtain “chance level” sensitivity. A Wilcoxon rank-sum test was used to compare the distributions of d’ values in each bin.

### Decoding Stimulus location

We decoded the information carried by a neural ensemble using a linear decoder. We used a L2-regularized logistic regression (LR) machine (Backen et al., 2018) to decode the attended location from increasingly larger ensembles. The details of the ensemble building procedure have been described elsewhere (Backen et al., 2018; Duong et al., 2019; Leavitt et al., 2017b, 2017a). Briefly, we started our ensemble of n =1 with the most informative single unit (based on all units’ individual decoding performance) and then iteratively paired it with all remaining units until we identified the unit that maximized the decoder’s performance and then added it to the ensemble (n = 2). Starting again with the best pair, we identified the best trio by repeating the same procedure, and the best quartet, and so forth. We repeated this procedure 100 times.

We normalized the firing rate of each unit across all trials by subtracting the mean and dividing by the standard deviation (z-score). These two parameters (mean and standard deviation) were estimated from the training set and applied to both the training set and the testing set. To assess the accuracy of the decoder, we used a cross-validation technique: The decoder was trained on 80% of the trials and then tested on the remaining 20% of trials (5-fold cross-validation). The LR was iteratively trained and tested on different subsamples of the trials until each trial was included in the training set at least once. Furthermore, we balanced the number of trials between the conditions by determining the sample size. Shuffled decoding performance (i.e., chance level) was obtained by randomizing the trial order 100 times. When testing the LR on such trials, we obtained the chance performance, which approximates 100*(1/N): N being the number of possible labels (e.g. N=2).

We integrated the firing rate of each unit within the whole population using several time windows centered at different epochs. We chose a 400 ms time window centered at cue epoch because it yielded good decoding accuracy. Therefore, we computed ensemble decoding accuracy over 400 ms windows centered at cue epoch for different ensemble sizes. We investigated whether the maximum decoding performance and corresponding ensemble size differed between conditions by comparing the maximum accuracy and corresponding ensemble size.

We also computed a mean decoding accuracy for the ensemble of neurons with the maximum decoding accuracy, the optimized ensemble, using a 400 ms window sliding by 100 ms over the time course of the experiment for each condition.

### Correlations

We computed noise correlations (r_noise_) by z-scoring the firing rate of each neuron for each unique condition, thus removing any possible changes in firing rate due to different responses to different stimuli. We then computed pairwise Pearson’s correlation coefficients between the z-scored spike counts.

We assessed the impact of correlations between pairs of neurons on optimized ensemble decoding accuracy by removing noise correlation between pairs. To destroy the noise correlation structure, we applied a randomizing procedure in which the trial order in each target location (e.g. target in left) was shuffled for each neuron individually, thus breaking up the simultaneity of recordings. This shuffling procedure was repeated 100 times. The difference between decoding performance with and without correlations was assessed using a paired sample Student’s t-test (p< 0.05).

## RESULTS

### Behavioral Performance

We trained two adult monkeys (“R” and “S”) to hold gaze on a central fixation point while covertly attending to one of two peripherally presented moving random dot patterns (RDPs). The dots in both RDPs always moved in the same directions but after a variable initial fixation period, they changed colors (e.g., in one RDP to red and in the other to green). The monkeys had to identify one of the two colored RDPs (the target) and attend to it while ignoring the other (the distractor) based on the color rule (Lennert & Martinez-Trujillo, 2011, 2013). Briefly, we arranged six iso-luminant colors in an ordinal hierarchy and the monkeys always had to attend to the stimulus with the higher-ranking color. After 706 ms, the colors of the RDPs returned to white. The animals were rewarded for releasing a lever after correctly detecting a brief motion direction change in the target (red in **Figure 1A, B**). In 50% of the trials, the distractor stimulus changed direction before the target. The animals had to ignore the distracter change and wait for the target change. In one set of trials the two stimuli were either presented bilaterally with equal probability of the target being on the left or right side of the fixation point (across condition, **Fig. 1A**). In another set of trials the stimuli were both presented on the same hemifield in the upper and lower quadrants (within left hemifield condition or within right hemifield condition) (**Fig. 1B**).

We compared the animals’ correct, false, and miss rates between the three conditions (**Fig. 1C-D**). There was a significant decrease in the proportion of correct trials in both monkeys in the within relative to the across condition ( *χ*^2^test p< 10^-3^). When examining the kind of errors the animals made, the proportion of responses to the distractor (i.e., false alarms) and the proportion of misses (i.e., no responses to a change in the target), increase significantly in the within relative to the across conditions in both monkeys ( *χ*^2^ test p< 10^-1^). In addition, there is no significant behavioral performance change between “within left” and “within right” conditions.

Consistent with previous studies (Carrasco et al., 2006; Rodriguez et al., 2010), these results indicate that attentional filtering is less effective when the target and distractor are in the same hemifield as compared to when they are across hemifields. The performance of animal S in the within condition was not different from chance. This may be attributed to a lesion related to the array implant or other unknown causes. Because we had no confidence the animal tried to perform the task in the within condition, data from this animal are not included in the rest of this paper. We will only examine neural data from monkey R.

### Single neuron analysis

We recorded the activity of single neurons using a 10×10 multielectrode array (**Fig. 2A**, Utah array (Normann & Fernandez, 2016)) and examined whether the units’ response was modulated by the location of the target stimulus using a one-way ANOVA (see methods). Some units responded more strongly when the target was on the left side of fixation point (ipsilateral to recording side) in across condition (example unit in **Fig. 2B**, paired t test p<10^-10^ on the average responses in the post cue 750 ms interval). Other neurons responded more strongly when the target was presented on the left lower quadrant in the within condition (example unit in **Fig. 2C**, paired t test p<10^-6^). The differences in response appear larger in the across condition than in the within condition indicating a stronger response modulation when the attended/unattended stimuli were in opposite hemifields.

To examine whether this pattern generalizes to the rest of the units we compared the proportions of location-selective units for four different task epochs in the across (n=525 units over 4 sessions), within left, and within right (n=331 units over 4 sessions) conditions. We found a significant increase of proportion of tuned units in cue and post-cue periods compared to the fixation period for three conditions (χ^2^-test, p < 10^-3^ for the across and within left condition and p < 0.05 for within right condition) (**Fig. 2D**, see insets for analysis periods). The proportion of location-selective units was significantly higher in the across relative to the within conditions in the cue and post cue periods (**Fig. 2E**, χ^2^-test, p < 10^-3^ for all comparisons). These data indicate that there were a higher number of neurons selective for the attended location in the across than in the within condition.

Next, we examine whether from one session to the next, there were differences between the proportion of tuned units over epochs. Overall, the proportion of units tuned for the different task epochs appears similar across recording sessions of the array indicating that the selectivity was stable over recording sessions (**Fig. 2F**). Together, our results indicate that single units in LPFC are significantly less selective for the attended location in the within relative to the across condition and that the proportion of selective units for different recording sessions was relatively stable. We cannot fully rule out that units were the same or different across rcoding sessions. However, a separate analyses of the recordings indicated that only 11% of the units remain stable from session to another using the MEA setup we have used here (Tremblay et al., 2015).

### Single Unit Discriminability

We assess whether the population of single neurons could discriminate between targets and distractors in the different conditions. We computed a measure of discriminability based on firing rates d-prime (d’ see methods) for the location-selective units (tuned units, 35% (n=183) across, 20% (n=66) within left and 6% (n=19)) in 100 ms bins with 1 ms sliding windows. To obtain an estimate of the discriminability predicted by chance we shuffled trial labels (50 shuffles) for each neuron, in all experimental conditions (Dashed line in **Fig. 4**). We conducted a Wilcoxon test for each pair of conditions for each time bin. All d’ were higher than predicted by chance during the cue and post cue period (Wilcoxon rank-sum test, p < 0.05) except for the within right condition. d’ was significantly higher in the across than in the within conditions starting from color cue onset and remaining until the end of the postcue period (Wilcoxon rank-sum test, p < 0.05). Overall, these results indicate that neurons in the LPFC better discriminate targets from distractors in the across relative to the within condition. For the case of the ‘within right’ condition discriminability was not higher than chance. The latter result may have been influenced by the smaller sample size in that condition.

**Figure 4:**
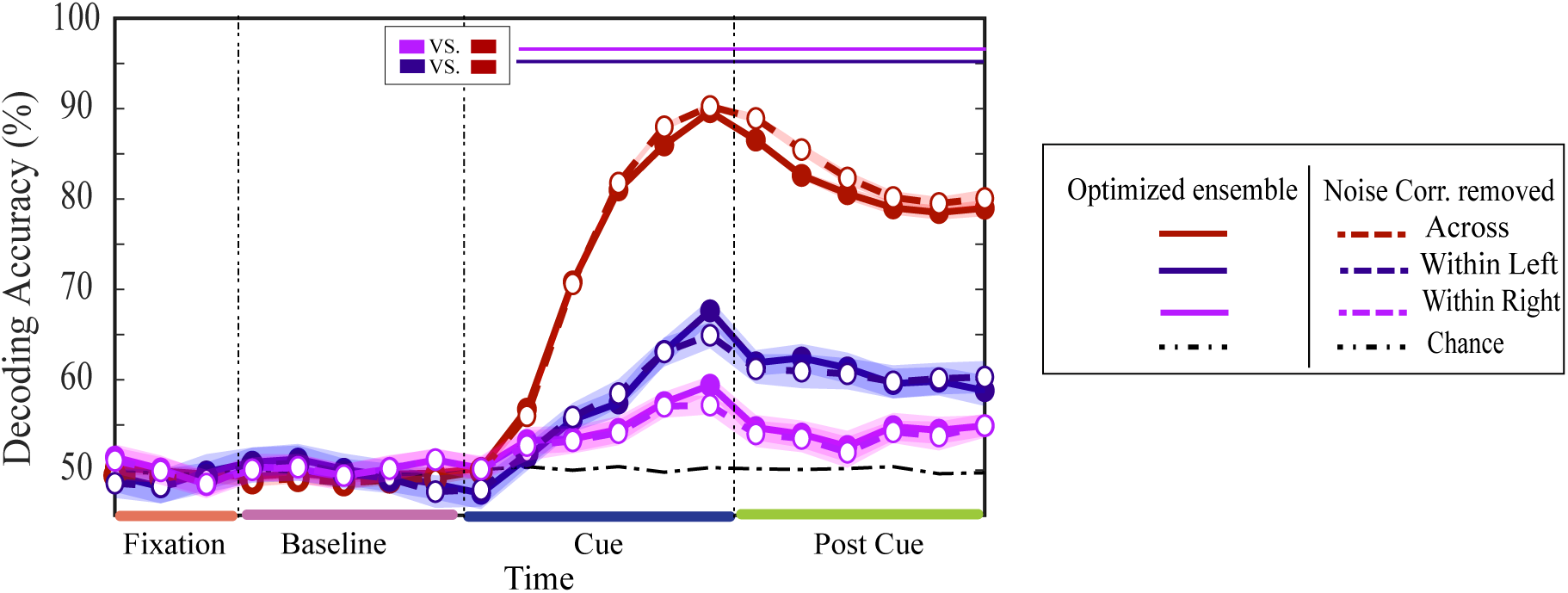
Effects of Correlations on Information Coding. Decoding accuracy using 400 ms sliding window over 100 ms intervals for optimized ensemble in the across (red lines with close circles), within left (blue lines with close circles) and within right (magenta lines with close circles) conditions. Dashed lines with open circles represent decoding accuracy without noise correlations. Black dashed dot line represents chance performance. Shading indicates SEM (±) for each time point. The straight lines at the top indicate statistically significant (Wilcoxon rank sum p< 0.05) differences between optimized ensemble performance in within left and within right conditions versus across condition.

### Performance of Neuronal Ensembles

Single unit analyses inform us about how a population of independently firing neurons encodes information about a stimulus or behavioral state. However, it has been shown that the performance of neuronal ensembles measured using classifiers can be different from that of single neurons and takes into account the correlation structure of neuronal ensembles (Backen et al., 2018; Leavitt et al., 2017b; Roussy, Luna, et al., 2021; Tremblay et al., 2015). We used a binary, linear classifier – the L2 regularized logistic regression machine (LR)– to decode target location in three conditions from the activity of the simultaneously recorded ensembles in correct trials. We assessed the decoder’s accuracy using a cross-validation technique in which it iteratively learned the neural response patterns to the stimulus conditions in 80% of the trials and was then tested on the remaining 20%.

We used running time windows of 400ms to compute firing rates (Backen et al., 2018). We progressively built neuronal ensembles (best ensemble) by starting with the most informative single unit and then iteratively adding units that maximized the decoder’s performance (see Leavitt et al. 2017 for methods). We also computed chance performance after randomizing the trial labels in the best ensembles and thus breaking up the simultaneity of the recordings and the shared trial-by-trial variability in the responses (i.e., noise correlations) (black dashed line in **Fig. 4**). Ensemble decoding accuracy was higher for the across condition (red line in **Fig. 4**) than for the within left (blue line) and within right (magenta line) conditions (paired t test, p < 10^-4^). In all conditions ensemble performance was significantly higher than chance level (Wilcoxon rank-sum test, p < 0.05). Interestingly, decoding accuracy was higher for ipsilaterally presented stimuli (within left) relative to the hemisphere where the array was implanted than for contralaterally presented stimuli (within right). However, this was not statistically significant (two-sample *t*-test, p ∼ 0.119).

We finally explored the effect of noise correlations on the decoding performance of the neuronal ensembles. We removed noise correlations from the optimized ensemble by shuffling the identity of trials belonging to the same trial condition (e.g. target is in the left) for each individual neuron (Tremblay et al., 2015). We found that reducing noise correlation did not significantly change decoding accuracy (**Fig. 4** dashed lines) (p >0.1 two sample t-test for all comparisons). This is concordant with reports of a complex relationship between noise correlations and information encoded in neurons populations (Leavitt et al., 2017b; Nogueira et al., 2019, 2020; Roussy, Luna, et al., 2021).

## DISCUSSION

We investigated the behavioral and neural correlates of attentional filtering when two stimuli (target and distractor) are presented within the same hemifield as compared to when they are across hemifields. Consistent with previous studies (Alvarez & Cavanagh, 2005; Awh & Pashler, 2000; Cavanagh & Alvarez, 2005; Chakravarthi & Cavanagh, 2009; Delvenne & Holt, 2012; Störmer et al., 2014), we found that behavioral performance in a spatial attention task is significantly decreased when stimuli are presented within the same visual hemifield. Moreover, we found that this behavioral effect is matched by corresponding correlates in the ability of LPFC single neurons and ensembles to discriminate between attended targets and ignored distractors. We found that in the within relative to the across condition there was: 1) a decrease in the proportion of attention selective units, 2) a decrease in d’ of single neurons, and 3) a reduced decoding performance of neuronal ensembles to discriminate the allocation of attention.

### Filtering Distractors Within versus Across Hemifields

Psychophysical research suggests that attentional resources are hemispheric-specific independent pools (Alvarez & Cavanagh, 2005; Delvenne & Holt, 2012; Walter et al., 2016). Basically, separate attentional resources exist for each hemifield. Our data shows that attentional filtering in LPFC neurons is diminished when monkeys allocated visuospatial attention to one of two stimuli when they were presented in the same hemifield relative to when they were presented in different hemifields. A study in macaques demonstrated that information about an object’s identity encoded by single neurons in LIP and LPFC was reduced when another object was in the same as compared to the opposite visual field (Buschman et al., 2011). Another single cell study in macaques demonstrated that when animals remember the location of two stimuli the responses of LPFC neurons decrease relative to when they remember the location of one stimulus (i.e., effect of memory load). Interestingly, this effect was much larger when the two stimuli were in the same hemifield that when they were in opposite hemifields and has been interpreted as evidence for independent working memory resources in each hemifield (Matsushima & Tanaka, 2014).

Our study design was different from the one of the previous studies in that we did not use a working memory paradigm in which stimuli had to be remembered. In our task the animals must use sustained attention to monitor a change in the target while ignoring the distracter so there is no need to use working memory. We have previously demonstrated that different subpopulations of neurons within the LPFC are activated by stimuli that are attended and remain visible (visual attention), and by stimuli that disappeared from the field of view and must be remembered (working memory) (Mendoza-Halliday & Martinez-Trujillo, 2017). However, the similarity between our findings and those reported by studies of working memory indicate that both cognitive functions may share similar hemifield-dependent constraints (Roussy, Mendoza-Halliday, et al., 2021). A possibility is that the results observed in working memory studies are due to attentional capacity limitations. Here one may consider that models of working memory include an attentional control system that direct attention to internal representations (Baddeley, 2010). Some studies have pointed out an overlapping between populations of neurons encoding working memory and attention (Lebedev et al., 2004; Mendoza-Halliday & Martinez-Trujillo, 2017; Panichello & Buschman, 2021; Roussy, Mendoza-Halliday, et al., 2021). In our view, attention and working memory are different functions in the sense one can attend without remembering it (Mendoza et al., 2011). The reverse may not be possible, one may not be able to effectively employ working memory without attention (Atkinson et al., 2018; Baddeley, 2010; Hitch et al., 2020). Practically, in many instances both functions may overlap (Roussy, Mendoza-Halliday, et al., 2021).

Our results may be related to the fact that stimuli across hemifields activate subpopulations of neurons including a single stimulus in the receptive field (RF) (Bullock et al., 2017). On the other hand, stimuli within the same hemifield are more likely to activate subpopulations including both stimuli in the RF. The former scenario may not trigger competitive interactions between neurons activated by the different stimuli. Such interactions occur via normalization mechanisms (Reynolds et al., 1999; Reynolds & Heeger, 2009) that decrease the firing rate to the attended stimulus due to activation of the normalization pool by the presence of the distractor. Decreases in firing rate leads to a deterioration in the representational capacity of populations encoding target features (McAdams & Maunsell, 1999). Within this framework is the lack of competitive interactions between neurons what allows better performance in the across hemifields condition.

In other words, independent stimulus representations may facilitate for the attentional system to dedicate separate processing resources to each stimulus, and animals may be able to attend to a target with minimal interference from the distracter. In agreement with this hypothesis, some studies have reported that although LPFC has a representation of stimuli on both visual hemifields, it has a bias for processing contralateral stimuli (Bullock et al., 2017; Lennert & Martinez-Trujillo, 2011, 2013). If such a bias exists for both hemifields, it would mean each hemisphere would have a non-overlapping representation of the contralateral visual hemifield. A distracter in the opposite hemifield may not ‘compete’ against an independent target representation to the same degree than a distracter within the same hemifield.

One may also argue that in early visual areas such as MT, with a contralateral representation of the visual field (Martinez-Trujillo & Treue, 2004; Niebergall et al., 2011; Yoo et al., 2018), competitive interactions only occur when stimuli are within the same hemifield. Indeed although MT receptive fields can be shaped by attention (Niebergall et al., 2011; Womelsdorf et al., 2008), they rarely cross the midline (Born & Bradley, 2005; Martinez-Trujillo & Treue, 2004; Treue & Martinez-Trujillo, 1999; Yoo et al., 2018). Interhemispheric effects have been rarely reported within this area or are small in magnitude and likely to occur via top-down bias through feedback projections (Bichot et al., 2015; Buffalo et al., 2010; Maunsell & Treue, 2006; Mendoza-Halliday et al., 2014; Moore & Armstrong, 2003; Moore & Fallah, 2004; Noudoost et al., 2010; Yoo et al., 2018; Zhou & Desimone, 2011). Thus, in early visual areas processing of stimuli in different hemifields occurs quasi-independently and therefore interactions within an area microcircuit between interhemifield representations are unlikely to occur.

### Decoding from Neuronal Ensembles

Using our ensemble building approach, we were able to decode the location of the target stimulus reliably above chance (Backen et al., 2018; Duong et al., 2019; Leavitt et al., 2017a, 2017b; Tremblay et al., 2015). The decoding of neuronal ensembles was higher in the across relative to the within condition, which agrees with the results of the single neuron analyses. Even for ensembles of the same size, decoding was always higher in the across relative to the within condition, therefore the differences in performance were not due to differences in the number of features (neurons) used by the classifier. Interestingly, removing noise correlations did not significantly change the decoding performance for any of the conditions. Nosie correlations in LPFC depend on the distance between neurons and similarity in their preferred feature (Leavitt et al., 2013). The latter result agrees with the complex effect of noise correlation on information decoding in neuronal ensembles (Averbeck et al., 2006; Leavitt et al., 2017a, 2017b; Moreno-Bote et al., 2014; Nogueira et al., 2019, 2020; Roussy, Luna, et al., 2021). Thus, at least in our data noise correlations may have not played a significant role in the bilateral field advantage effect.

### Conclusions

Our results show a bilateral field advantage of spatial attention at the level of behavior. We also report a neural correlate of this effect at the level of single neurons and neuronal ensembles in the LPFC of macaque monkeys. Our findings suggest that the bilateral field advantage occurs due to the decreased competitive interaction between population of neurons representing stimuli falling across different hemifields. We propose this effect may emerge in the LPFC, but it may get magnified by the lack of local interactions occurring in early visual areas between neurons with a single hemifield representation when attention operates across hemifields. One could interpret our results as existence of separate processing resources for different hemifields representations during the allocation of attention.

## ACKNOWLEDGEMENTS

We thank Mr. Stephen Nuara for animal assistance and Mr. Walter Kucharski for technical support. We also thank Mr. Rishi Rajalingham for valuable comments on a previous version of this manuscript.

